# Macrophage internalization creates a multidrug-tolerant fungal persister population, providing a permissive reservoir for the emergence of drug resistance

**DOI:** 10.1101/2022.10.21.513290

**Authors:** Amir Arastehfar, Farnaz Daneshnia, Nathaly Cabrera, Suyapa Penalva-Lopez, Jansy Sarathy, Matthew Zimmerman, Erika Shor, David S. Perlin

## Abstract

*Candida glabrata* is a major fungal pathogen notable for causing recalcitrant infections, rapid emergence of drug-resistant strains, and its ability to survive and proliferate within macrophages. Resembling bacterial persisters, a subset of genetically drug-susceptible *C. glabrata* cells can survive lethal exposure to the fungicidal echinocandin drugs. Herein, we show that macrophage internalization induces cidal drug tolerance in *C. glabrata*, expanding the persister reservoir from which echinocandin-resistant mutants emerge. We show that this drug tolerance is associated with non-proliferation and is triggered by macrophage-induced oxidative stress, and that deletion of genes involved in reactive oxygen species detoxification significantly increases the emergence of echinocandin-resistant mutants. Finally, we show that the fungicidal drug amphotericin B can kill intracellular *C. glabrata* echinocandin persisters, reducing emergence of resistance. Our study supports the hypothesis that intra-macrophage *C. glabrata* is a reservoir of recalcitrant/drug-resistant infections, and that drug alternating strategies can be developed to eliminate this reservoir.

## Introduction

The yeast *Candida glabrata* is a component of the human microbiome that inhabits a wide range of mucosal surfaces and is a prevalent opportunistic fungal pathogen causing bloodstream infections around the world (1–5). A prominent increase in azole resistant *Candida* species, including *C. glabrata*, in recent years has promoted the use fungicidal (cidal) echinocandin drugs, such as caspofungin and micafungin, which interfere with cell wall biosynthesis, as prophylaxis and first line therapy against candidemia (6, 7). In turn, this change in practice has been accompanied by high rates of echinocandin- and multi-drug resistant *C. glabrata* isolates in numerous clinical centers (6). This worrying spike in the number of drug resistant *C. glabrata* isolates, combined with the cytotoxicity of polyenes, such as amphotericin B, has greatly limited therapeutic options to treat candidemia (8).

Numerous studies have documented that echinocandin-resistant (ECR) *C. glabrata* isolates emerge from genetically related susceptible cells during the course of infection (9–14). It has been hypothesized that a subpopulation of echinocandin-susceptible *C. glabrata* cells survives fungicidal concentrations of echinocandins in the host and this leads to the emergence of ECR isolates. In bacteriology, such cells, termed “persisters”, have been well-studied (16, 17). In planktonic cultures, bacterial persisters are defined by their biphasic killing dynamic, where a growth-restricted subpopulation can tolerate and survive high concentrations of cidal antibiotics lethal to their clonal susceptible kin. Persisters can be induced by environmental stresses, such as starvation, low pH, and reactive oxygen species (ROS) (16–24). Thus, intracellular bacteria, such as *Mycobacetrium tuberculosis, Listeria monocytogenes* and *Staphylococcus aureus*, have significantly increased antibiotic tolerance compared to counterparts grown in culture due to stresses imposed by host cells (16–24). Unlike antibiotic-resistant cells, persisters do not carry heritable mutations, and their progeny retain antibiotic susceptibility. After the cessation of antibiotic treatment, such persisters can reinitiate growth, causing a relapse of infection. Importantly, persisters can acquire the genetic mutations leading to mechanism-specific antibiotic resistance (16, 17).

In mycology, it is assumed that persisters are formed predominantly in biofilms following exposure to fungicidal drugs, but rarely under planktonic conditions (25). Interestingly, it has also been suggested that *C. glabrata* isolates poorly produce persister cells even in the context of biofilms (25–27). Our group, however, has characterized a small subpopulation of *C. glabrata* persisters *in vitro* (which we referred to as “drug-tolerant cells”) surviving supra-high MIC concentration of echinocandins (15, 28). Such subpopulations growing or surviving supra-high MIC concentrations of static or cidal antifungal drugs, respectively, have been observed in multiple fungal species and proposed to be involved in therapeutic failure, emergence of resistance, and poor clinical outcomes (29–32).

A unique feature of *C. glabrata* is that it can survive and replicate inside macrophages (4). Yet, it has not been determined how intracellular *C. glabrata* (ICG) responds to antifungal drugs, particularly the cidal echinocandins, and whether macrophages may represent a persister reservoir in the context of infection. Here, we show that internalization by macrophages significantly increases the abundance of multidrug-tolerant *C. glabrata* persister cells resulting in a significantly higher rate of ECR colonies. Interestingly, isolates lacking genes involved in ROS detoxification had greatly increased formation of ECR mutants, pointing to ROS as a potential inducer of mutagenesis. Finally, amphotericin B, a fungicidal drug with a different mode of action, was able to kill *C. glabrata* persisters induced by micafungin and significantly reduce ECR colony formation. Altogether, this study demonstrates that *C. glabrata* harbored within host macrophages during infection constitutes a persister reservoir and a source of ECR isolates, and points towards strategies to eliminate persisters and reduce emergence of drug-resistant strains.

## Results

### ICG shows a persister phenotype

To assess the impact of antifungal drugs on ICG, we selected two isolates with similar MIC values: reference strain CBS138 (sequence type 15, ST15) and strain Q36, a clinical isolate belonging to ST3. THP1 macrophages exposed to *C. glabrata* cells for 3 hours were extensively washed to remove non-adherent yeast cells and treated with RPMI containing 2X MIC of the fungistatic azoles fluconazole and voriconazole, and fungicidal echinocandins micafungin, and polyene amphotericin B. The two strains were also incubated in RPMI medium alone containing the same drug concentrations. The survival/proliferation of both ICG and planktonic cells were measured by colony forming unit (CFU) counts and normalized against respective untreated controls at 3, 6, and 24 hours post treatment (pst). Interestingly, drug-treated ICG showed significantly higher survival compared to drug-treated planktonic cells (Figure 1a). Next, we treated ICG and planktonic cells with a wide range of micafungin concentrations (2X-256X MIC) and observed that the survival of ICG was 100-1000-fold higher than that of planktonic cells, especially at 24 hours pst (Figure 1b). This high survival rate of ICG was reminiscent of intracellular bacterial persister cells (16, 17). To ask whether ICG fulfill other criteria for being considered persisters, we selected a single micafungin concentration (0.125μg/ml, 8X MIC) and investigated multiple parameters used to specify bacterial persisters.

**Figure 1.**
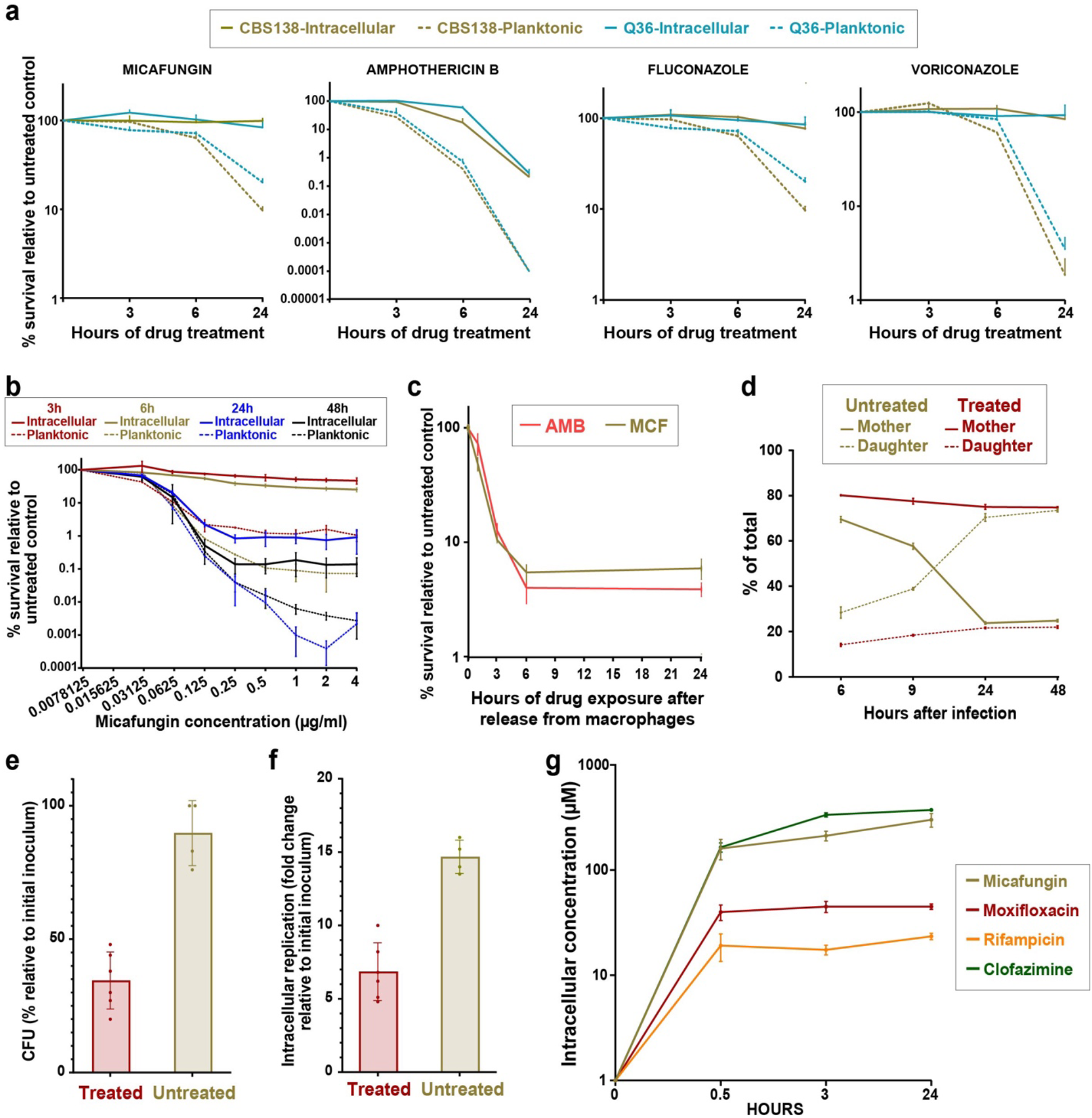
Intracellular *C. glabrata* (ICG) cells show multiple characteristics of persisters. **a.** Survival rates of ICG exposed to micafungin (0.03μg/ml), AMB (1μg/ml), fluconazole (8μg/ml), and voriconazole (0.06μg/ml) were significantly higher than those of their planktonic counterparts. **b.** ICG survival was higher than that of planktonic cells over a range of micafungin concentrations **c.** ICG exposed to 8X MIC of micafungin (0.125μg/ml) or AMB (4μg/ml) show a biphasic killing pattern. **d.** Proportions of daughter (AF647 ConA-positive) and mother cells (FITC- and AF-647 ConA-positive) in micafungin-treated ICG remain stable over the course of treatment, indicating a lack of proliferation, whereas untreated ICG show a marked increase in the proportion of daughter cells. **e.** Propidium iodide (PI)-negative *C. glabrata* cells released from micafungin-treated macrophages have lower culturability, as assessed by colony forming units (CFU), than PI-negative *C. glabrata* released from untreated macrophages. **f.** PI-negative *C. glabrata* released from micafungin-treated macrophages are capable of reinfecting new macrophages, albeit at a lower rate relative to untreated counterparts. **g.** Micafungin penetrated macrophages and accumulated there at a high concentration, similar to clofazimine and higher than moxifloxacin and rifampicin. PI: propidium iodide, AF-647 ConA: Alexa Flour 647-conjugated concanavalin A, FITC: fluorescein isothiocyanate, ICG: intracellular *C. glabrata*, MCF: micafungin AMB: amphotericin B, MIC: minimum inhibitory concentration, CFU: colony forming units, LC-MS/MS: liquid chromatography-tandem mass spectrometry.

First, to ask whether ICG show the hallmark persister biphasic killing, THP1 macrophages were infected with *C. glabrata* and incubated for 3 hours. Subsequently, nonadherent yeast cells were removed by extensive washing and survival was assessed at 1-, 3-, 6-, and 24-hours pst. Indeed, the ICG showed the typical biphasic killing to both micafungin and amphotericin B (Figure 1c).

To test whether ICG cells are nonproliferating, prior to infecting macrophages we stained the initial inoculum with isothiocyanate (FITC), which does not transfer to daughter cells. At designated timepoints, the ICG was counterstained with Alexa Flour-647 (AF647), which stains all cells (mothers and daughters), and the micafungin-treated ICG and untreated controls were released from macrophages and subjected to flow cytometry. *C. glabrata* cells single positive for AF647 represented daughter cells and therefore indicated proliferation within macrophages, whereas cells doubly stained with both FITC and AF647 represented the mother cells. Interestingly, while untreated ICG showed a dramatic expansion of daughter cells, upon micafungin treatment the proportion of mother and daughter cells did not vary at 3, 6, 24, or 48 hours of micafungin exposure (Figure 1d), indicating low or no proliferation.

Next, we tested whether after a 24-hour exposure to micafungin ICG had acquired genetic determinants of echinocandin resistance (mutations in the hot-spot (HS) regions of *FKS1* and *FKS2*). Treated ICG was plated on YPD containing 0.125μg/ml micafungin to capture the ECR colonies. We selected this concentration because our pilot studies had shown that it could detect various *FKS* HS mutations after plating a range of cell numbers (10-10^6^ cells). We found that ICG did not produce any ECR colonies, consistent with a lack of *fks* mutations in ICG, and that colonies obtained from them showed killing dynamics similar to the parental strain.

Finally, we asked whether *C. glabrata* released from micafungin-treated macrophages are culturable and whether they are capable of initiating another cycle of infection and proliferation inside macrophages. After a 24-hour micafungin treatment, *C. glabrata* cells were released from macrophages and stained with propidium iodide (PI). PI-negative (PI-) cells were selected and either plated on YPD plates or used to infect macrophages, and proliferation within macrophages was assessed after 72 hours. We found that *C. glabrata* released from micafungin-treated macrophages were capable of forming colonies (Figure 1e) and reinfecting and proliferating within macrophages (Figure 1f), but at significantly lower rates than planktonic cells, which is similar to results obtained with bacterial persisters (33).

### Micafungin efficiently penetrates into macrophages and rapidly reaches supra-MIC concentrations

We sought to determine the dynamics of micafungin penetration into the macrophages to assess if the high survival of ICG cells was simply due to poor micafungin penetration. We directly measured the concentration of micafungin inside THP1 macrophages cultured in 4 μg/ml (3.15 μM) of the drug using liquid chromatography with tandem mass spectrometry (LC/MS/MS), which can accurately predict drug penetration at the site of infection *in vivo* (34, 35). We measured the intracellular (IC) and extracellular (EC) concentrations of micafungin at 30 minutes and 3 and 24 hours pst. We included three control drugs with well-established macrophage penetration capacities, each set at 5 μM: rifampicin (poorly penetrating), moxifloxacin (moderately penetrating), and clofazimine (highly penetrating). As expected and consistent with previous studies (35–37), rifampicin showed the poorest penetration, whereas moxifloxacin and clofazimine moderately and highly penetrated the macrophages, respectively (Figure 1g). Micafungin showed comparable penetration into the macrophages to clofazimine despite being used at a lower concentration. This result was consistent with previously reported strong penetration and accumulation of echinocandins in macrophages in clinical settings (38–40) and indicated that ICG retain high viability even though the intracellular micafungin concentration reaches >100 μM, or 256X of the MIC, by 0.5 hour pst (Figure 1g).

### Internalization of *C. glabrata* cells by THP1 macrophages significantly increases their tolerance to cidal antifungals

Various environmental stresses are well known to induce cross-stress tolerance, including tolerance to antibiotics and antifungals (16–24). In particular, macrophage internalization increases the number of antibiotic persisters by exposing bacteria to an array of stresses (16–24). Accordingly, we tested whether internalization of *C. glabrata* by macrophages increased their cidal drug tolerance. *C. glabrata* cells were used to infect macrophages, cultured for 3, 6, 24, and 48 hours, then released from macrophages, exposed to micafungin (0.06 μg/ml) for one hour and plated on drug-free YPD for CFU counts. Control planktonic cells were cultured in RPMI but otherwise treated identically. Interestingly, ICG showed a significantly higher survival after treatment, especially at 3 hours, compared to planktonic cells (Figure 2a). Also, consistent with the observation that stationary phase bacteria are significantly enriched for persisters (41), we found that planktonic *C. glabrata* cells at 24 and especially 48 hours showed the highest survival rates upon micafungin exposure (Figure 2a).

**Figure 2.**
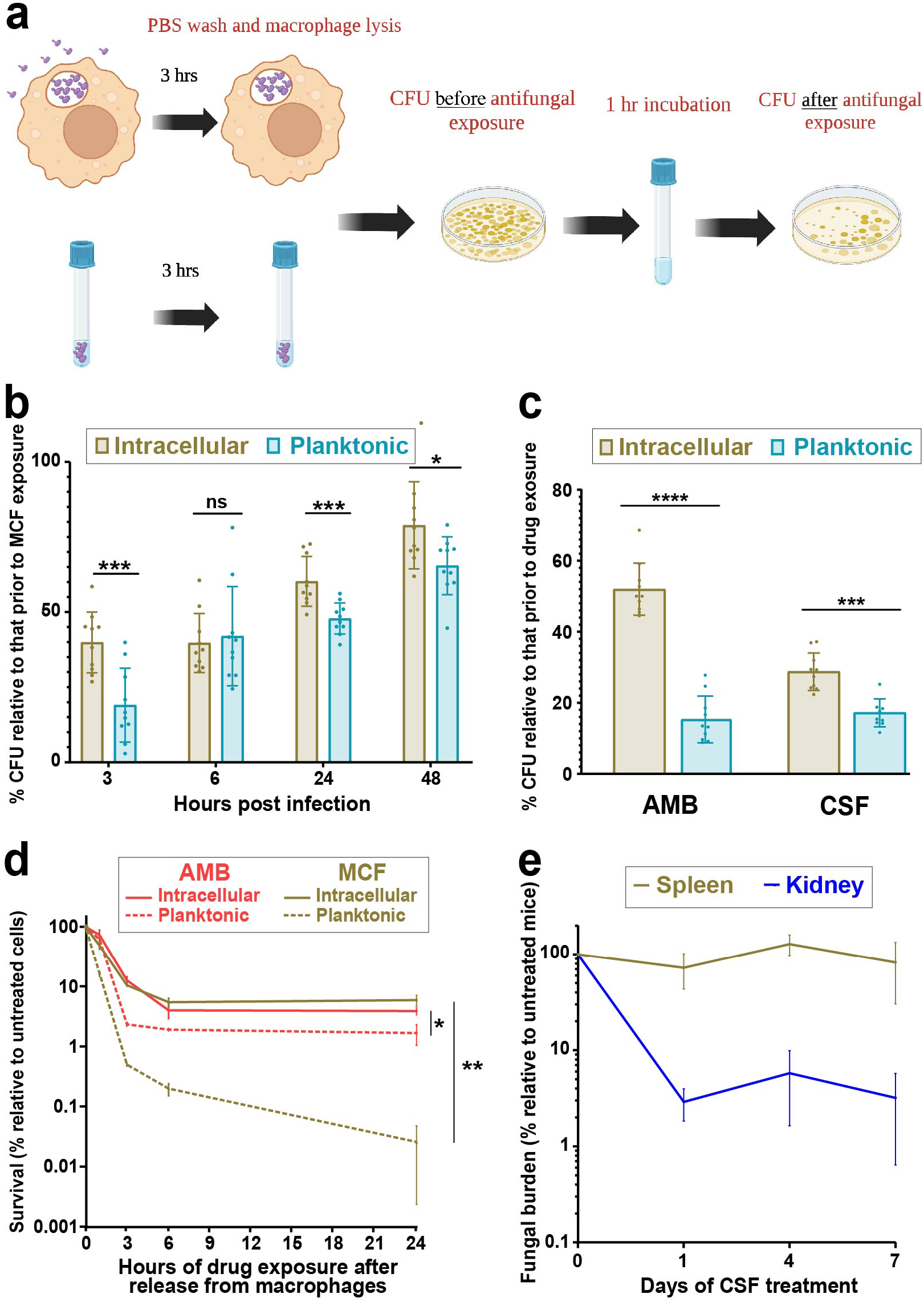
Internalization of *C. glabrata* cells by macrophages enhances their tolerance to cidal antifungal drugs. **a.** *C. glabrata* cells either incubated in RPMI or macrophages for 3 hours and their survival were assessed before and after one hour exposure to micafungin (0.06μg/ml). **b.** *C. glabrata* released from macrophages at different times post-infection showed significantly higher survival upon treatment with 0.125 μg/ml micafungin for 1 hour compared to similarly treated planktonic cells (* = p<0.03, ** = p<0.003, *** = p<0.0003, paired t-test). **c.** *C. glabrata* released from macrophages 3 hours post-infection and treated for 1 hour with either caspofungin (0.125μg/ml) or amphotericin B (2μg/ml) showed higher survival than similarly treated planktonic cells (** = p<0.003, *** = p<0.0003, paired t-test). **d.** *C. glabrata* cells released from macrophages 3 hours post-infection and treated with 0.125 μg/ml micafungin or 2 μg/ml amphotericin B showed significantly higher survival than similarly treated planktonic cells over a wide range of timepoints (* = p<0.03, paired t-test). **e.** *C. glabrata* burden in the spleen (a macrophage-rich organ) was not significantly reduced by treatment with a humanized dose of capsofungin (5 mg/kg), in contrast to the kidney. MCF: micafungin, AMB: amphotericin B, CSF: caspofungin, CFU: colony forming units.

To assess whether internalization by macrophages renders ICG more tolerant to other cidal antifungals, *C. glabrata* cells were used to infect macrophages or inoculate RPMI and 3 hours later exposed to 4X MIC of caspofungin or amphotericin B for one hour. Interestingly, ICG again showed a significantly higher survival in both caspofungin and amphotericin B relative to their planktonic counterparts (Figure 2b).

Finally, to understand how long the higher drug tolerance of ICG cells lasts, exponentially growing *C. glabrata* cells (CBS138) were either incubated in RPMI or internalized by macrophages for 3 hours, released, and then exposed to either micafungin (0.125μg/ml) or amphotericin B (4μg/ml) for 1, 3, 6, or 24 hours. Strikingly, the high tolerance level of ICG cells was detected up until 24 hours and was the highest at 24-hour time-point, especially for micafungin (Figure 2c). Altogether, these experiments showed that ICG cells display a long-lasting multidrug tolerant phenotype.

### *C. glabrata* cells harbored in the spleen have higher caspofungin tolerance than those harbored in the kidney

Because ICG showed significantly higher survival after release from macrophages, we asked whether during systemic infection *C. glabrata* harbored in the spleen, which contains one of the largest macrophage populations in the body (42), may have higher echinocandin tolerance than *C. glabrata* cells harbored in another organ. Mice were systemically infected with BYP40, a clinical *C. glabrata* strain, and either treated with a humanized dose of caspofungin (5mg/kg) every day following infection or received PBS alone (control). Spleen and kidney were harvested on days 1, 4, and 7, homogenized, and plated on YPD for CFU counts. In line with our hypothesis, we found that whereas kidney *C. glabrata* burdens significantly decreased in caspofungin-treated mice, spleen fungal burdens were unaltered by treatment (Figure 2d), indicating that macrophage-rich anatomical niches likely contain drug-tolerant persister *C. glabrata* cells.

### ROS produced by THP1 macrophages renders *C. glabrata* more tolerant to micafungin

To understand which stress applied by macrophages renders *C. glabrata* tolerant to cidal antifungal drugs, we carried out an *in vitro* experiment, in which *C. glabrata* cells were incubated in RPMI containing 10 mM H_2_O_2_ (representing ROS), pH5 RPMI (representing vacuolar acidity), spent RPMI (representing nutrient-poor vacuolar environment), spent RPMI+10 mM H_2_O_2_, or regular control RPMI. Next, at designated time-points the *C. glabrata* cells were exposed to micafungin (0.06 μg/ml) for one hour. Interestingly, *C. glabrata* cells incubated in either spent RPMI+H_2_O_2_ or H_2_O_2_ alone (and, to a lesser extent, those in spent RPMI) showed significantly increased tolerance to micafungin, whereas those incubated in acidic medium showed similar killing rates as controls (Figure 3a).

**Figure 3.**
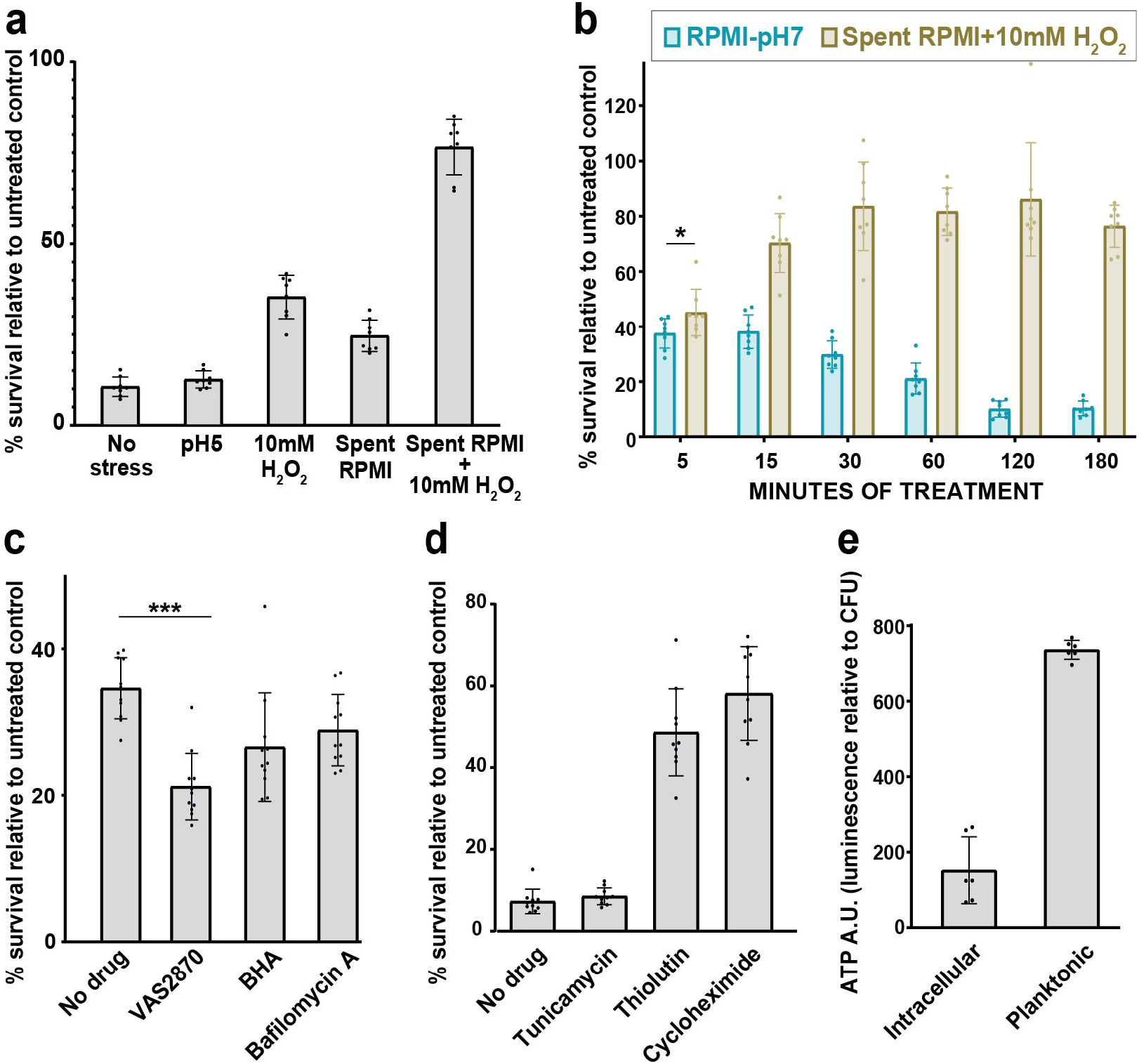
Oxidative stress and starvation drive *C. glabrata* into a drug-tolerant, persister-like state. **a.** Oxidative stress (RPMI+10mM H_2_O_2_), starvation (spent RPMI), and especially oxidative stress in combination with starvation (spent RPMI+10mM H_2_O_2_) increased the survival of *C. glabrata* after exposure to 0.125μg/ml micafungin, whereas low pH (RPMI pH5) had no effect. **b.** The effect of oxidative and starvation stresses on survival was evident as early as after 15 minutes of stress (* = p<0.03, ** = p<0.003, modified t-test). **c.** Inhibiting macrophage ROS production by using a NOX inhibitor (VAS2870) or an antioxidant (BHA) resulted in decreased survival of macrophage-released *C. glabrata* after 1 hour in 0.06 μg/ml micafungin, whereas treating macrophages with a vacuolar ATPase inhibitor (Bafilomycin A) had a weaker effect. **d.** *C. glabrata* cells incubated in RPMI containing sublethal concentrations of a transcription inhibitor (thiolutin 5μg/ml) or a translation inhibitor (cycloheximide 1μg/ml) had a significantly higher tolerance to micafungin (0.06 μg/ml) compared to *C. glabrata* treated with an endoplasmic reticulum stressor (tunicamycin 5μg/ml) or untreated cells. **e.** ATP levels of ICG as measured by a luciferase-based assay were significantly lower than in planktonic cells. ICG: intracellular *C. glabrata*, NOX: NADPH oxidase, BHA: Butylated droxyanisole, ROS: Reactive oxygen species, RPMI: Roswell Park Memorial Institute.

To determine how soon the combination of nutrient deprivation and ROS induces drug tolerance in *C. glabrata*, we incubated *C. glabrata* cells in spent RPMI+H_2_O_2_ for 5, 15, 30, 60, and 120 minutes, exposed the cells to 0.06 μg/ml micafungin for one hour, and assessed survival by CFU counts. Consistent with studies of bacterial persisters (20, 22, 24), we found that a high level of tolerance can be induced as early as after 15 minutes of incubation in spent RPMI+H_2_O_2_ (Figure 3b and Supplementary Figure 1).

To test directly whether macrophage-produced ROS enhances ICG drug tolerance, THP1 macrophages were pretreated with an antioxidant (butylated hydroxanisole, BHA) or a NAPH oxidase inhibitor (VAS2870) prior to *C. glabrata* infection (22). We also pretreated macrophages with bafilomycin A (Baf A), a vacuolar ATPase inhibitor that prevents phagosome acidification (43). Untreated macrophages served as controls. We found that ICG released from macrophages pretreated with BHA or VAS2870 showed significantly lower survival than ICG released from untreated macrophages (Figure 3c), indicating that macrophage ROS are a key inducer of ICG drug tolerance. Of note, ICG released from BafA-treated macrophages also had a lower survival rate than controls, which may indicate vacuole acidification may also partly contribute to the higher tolerance of ICG to micafungin.

It has been shown that *C. albicans* and *C. glabrata* cells internalized by macrophages for 3 hours suppress their transcription and translation machineries (44, 45). Thus, we tested whether inhibiting transcription or translation using sublethal concentrations of thiolutin and cycloheximide (4), respectively, could enhance the tolerance of *C. glabrata* cells to micafungin. Indeed, inhibition of either transcription or translation significantly increased *C. glabrata* micafungin tolerance (Figure 3d). We hypothesized that the observed macrophage-induced down-regulation of transcription and translation could be due to a reduction in ICG ATP stores, as it has been shown that macrophage-produced ROS impairs the TCA cycle, which leads to a sharp decrease in endogenous ATP levels (22, 23, 46). Indeed, we found that the ATP level of ICG cells was significantly lower than that of planktonic cells (Figure 3e). Together, these data indicate that macrophage engulfment exposes *C. glabrata* to environmental stresses (ROS, nutrient deprivation, and low pH), lowering its ATP levels and prompting it to downregulate transcription and translation and rapidly acquire a drug-tolerant state, leading to persistence.

### Internalization by macrophages significantly increases the emergence of ECR colonies

It has been shown that bacterial persisters constitute a reservoir from which drug resistant colonies can emerge (47), and the same has been hypothesized for fungal cells(15, 28). Therefore, we asked whether ICG is a source of ECR mutations. We infected macrophages with 5 clinical strains belonging to different STs as well as the reference strain CBS138 and monitored the emergence of ECR mutations in ICG in the presence of micafungin (0.125μg/ml). To detect ECR cells, *C. glabrata* were released from macrophages at 24-hour intervals and plated on micafungin-containing YPD plates (0.125 μg/ml). The initial inoculum of each isolate was also plated on YPD+micafungin to verify that it did not contain ECR cells. Of note, the majority of clinical ECR isolates associated with therapeutic failure harbor mutations in HS1 of *FKS2*, followed by HS1 of *FKS1* (48, 49). Thus, the ECR colonies growing on YPD+micafungin plates were subjected to sequencing of the four *FKS* HSs (*FKS1* HS1 and 2, *FKS2* HS1 and 2) (50). As in previous experiments (Figure 1a, Figure 3c), both ICG and planktonic *C. glabrata* showed declining survival over time up to 48 hours of micafungin treatment, with ICG being significantly more tolerant to micafungin than planktonic cells (Figure 4). After 48 hours, however, micafungin-treated planktonic cells displayed a CFU rebound (Figure 4), which is reminiscent of phenotypic resistance (17). Nevertheless, ICG produced a significantly higher frequency of ECR colonies compared to planktonic cells, and all these ECR colonies harbored well-known and clinically relevant mutations in HS1 of *FKS2* (Figure 4, Supplementary Table 1). This result indicates that macrophage-engulfed *C. glabrata* may constitute a reservoir from which ECR *C. glabrata* cells can emerge.

**Figure 4.**
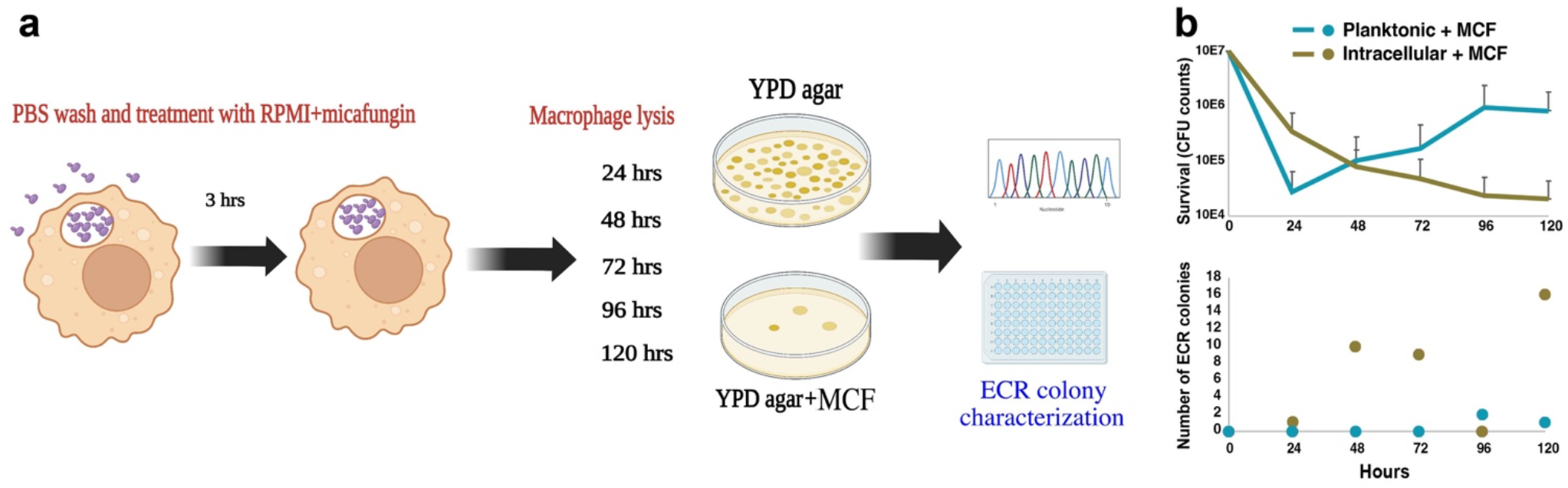
ICG develops echinocandin-resistant mutations. **a.** Echinocandin resistance and survival rates of *C. glabrata* cells incubated in macrophages and RPMI containing micafungin (0.125μg/ml) were evaluated. At each time-point, *C. glabrata* cells were collected, a portion was spread on YPD to assess survival and the other portion was plated on YPD plates containing micafungin (0.125μg/ml) to determine the ECR rate. **b.** The dynamics of killing (top panel) and the numbers of ECR colonies (bottom panel) detected at different time-points are shown for both ICG and planktonic cells cultured in the presence of 0.125 μg/ml micafungin. ICG: intracellular *C. glabrata*, ECR: echinocandin resistant.

### Deletion of ROS detoxifying genes markedly increases the frequency of ECR emergence

ROS are generally thought to be detrimental to cells because of their capacity to damage cellular macromolecules. However, both bacterial and fungal cells also use endogenously generated ROS to induce programmed genetic instability under specific circumstances (51, 52). Our group previously showed that although echinocandins induce endogenous ROS in *C. glabrata*, the use of ROS scavengers did not enhance the survival of *C. glabrata* in echinocandin presence and that, surprisingly, *C. glabrata* downregulated ROS detoxifying genes upon echinocandin exposure (28). To ask whether ROS are involved in persistence and emergence of ECR mutants, we deleted genes with well-known functions in ROS detoxification: catalase (*CAT1*, CAGL0K10868g), glutathione oxidoreductase (*GRX2*, CAGL0K05813g), manganese superoxide dismutase (*SOD2*, CAGL0E04356g), and three transcription regulators of oxidative stress responses, namely *SKN7* (CAGL0F09097g), *MSN4* (CAGL0M13189g), and *YAP1* (CAGL0H04631g) (28, 53). Additionally, because it has been shown that in *Mycobacetrium tuberculosis* isocitrate lyase mutants have elevated antibiotic-induced ROS levels (54), we also deleted *C. glabrata* isocitrate lyase gene *ICL1* (CAGL0L09273g).

All deletion mutants grew normally in YPD broth (Figure 5a). Consistent with their compromised ROS detoxification activities, most of the mutants showed increased ROS levels in the presence of micafungin (0.125μg/ml) (Figure 5b). To measure their sensitivity to exogenous ROS, we measured their survival in RPMI containing 10 mM of H_2_O_2_. Consistent with a previous study (53), we found that deleting *CTA1* and the two main transcription factors regulating its expression, *YAP1* and *SKN7*, significantly impaired strain survival in the presence of H_2_O_2_, whereas the other mutants were unaffected (Figure 5c). We also measured the mutants’ survival and proliferation within macrophages and found that *skn7*Δ, *sod2*Δ, and *yap1*Δ mutants showed significantly reduced survival 24 hours after infection, whereas only the *sod2*Δ mutant showed significantly reduced survival 48 hours after infection (*cta1*Δ and *icl1*Δ mutants did not reach the statistical significance threshold with p=0.06 for both) (Figure 5d). These results suggested that either the *C. glabrata* ROS detoxification pathways are redundant (55) or the ROS levels inside macrophages were not high enough to strongly impair the mutants.

**Figure 5.**
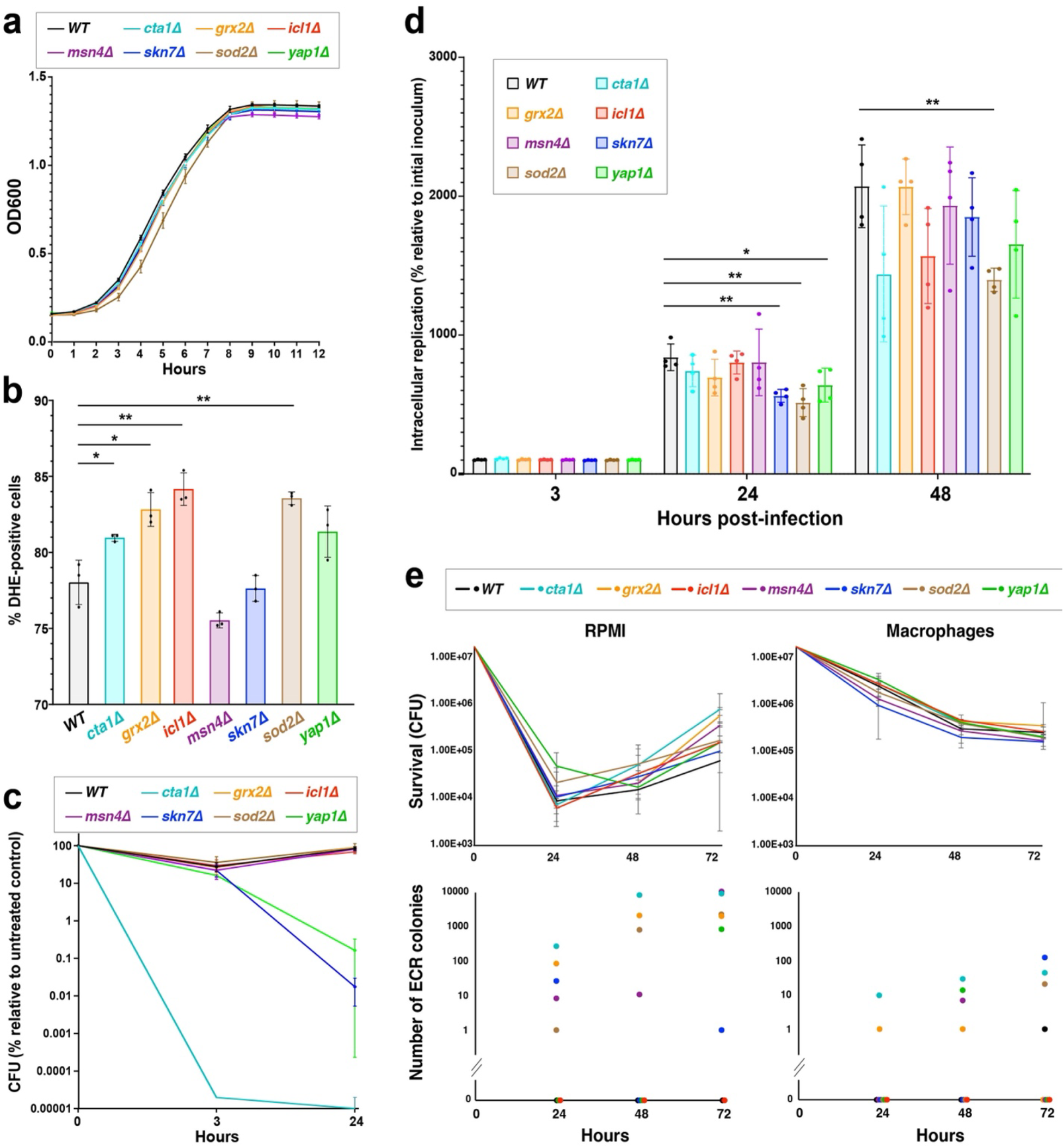
Deletion of *C. glabrata* ROS detoxification genes promotes the emergence of ECR mutants in both ICG and planktonic cells. **a.** ROS mutants had normal growth rates in YPD broth. **b.** With the exception of *msn4Δ* and *skn7Δ*, the ROS detoxification mutants had significantly higher superoxide levels compared to the WT strain when exposed to micafungin (0.125μg/ml), as measured by DHE straining intensity (* = p<0.03, ** = p<0.003, paired t-test). **c.** In the presence of 10mM H_2_O_2_ *ctalΔ, skn7Δ*, and *yaplΔ* mutants had significantly reduced survival compared to the WT strain. **d.** Several ROS mutants had mild survival/replication defects inside THP1 macrophages at 24-hour and 48-hour time-points (* = p<0.03, ** = p<0.003, modified t-test). **e.** The mutants were exposed to 0.125 μg/ml micafungin and survival and ECR colony formation were monitored. Although the mutants showed comparable survival in micafungin to the WT strain (top panel), they produced a significantly higher number of ECR colonies under either planktonic or intra-macrophage conditions (bottom panel). WT: wild type, YPD: yeast extract-peptone-dextrose, ICG: intracellular *C. glabrata*, DHE: dihydroethidium.

Next, we assessed the mutants’ survival in the presence of micafungin and the emergence of ECR mutations either when cultured in RPMI or harbored within macrophages. Interestingly, although none of the mutants were significantly impaired for survival in the presence of micafungin, both ICG and planktonic ROS detoxification mutants (except *icl1Δ*) showed greatly increased ECR colony emergence relative to the WT strain (Figures 5e and Supplementary Table 2). ECR frequencies were even higher in planktonic cells than in ICG, suggesting that it is not macrophage-produced exogenous ROS but *C. glabrata-produced* endogenous ROS may promote this mutagenesis. Interestingly, we did not recover any ECR colonies from the *icl1Δ* strain, although it had no obvious survival defect, despite repeating the experiment 3 times with a 120 hrs follow-up. This observation may point to the potential importance of *ICL1* for cellular processes linked to mutagenesis. Given the absence of this gene in human genome and that this gene is evolutionarily conserved across a wide range of pathogens (56), it may be a promising target to reduce emergence of ECR mutants in *C. glabrata*.

### Micafungin-tolerant ICG are effectively killed by amphotericin B

We have shown that the nutritional and ROS stresses imposed by macrophages on ICG drive the fungus into a non-proliferative, low ATP state that is highly tolerant to echinocandins. In principle, this persister population may be susceptible to a different class of drug whose cidal activity is not strongly affected by the metabolic or proliferative state of the pathogen. Indeed, in the bacterial field cidal antibiotics have been classified as either strongly dependent on metabolism (SDM) or weakly dependent on metabolism (WDM) (57–59). For *Candida* species, the metabolic dependency of cidal antifungal drugs has not been fully examined. Thus, we compared the metabolism dependencies of micafungin (0.125 μg/ml), caspofungin (0.25 μg/ml), and amphotericin B (2 μg/ml) against *C. glabrata* cells incubated in 100%, 20%, and 2% RPMI. Not surprisingly, micafungin and caspofungin were significantly affected by RPMI concentration, classifying them as SDM drugs, whereas the lethality of amphotericin B was much less dependent on RPMI concentration, classifying it as a WDM drug (Figure 6a). The strong dependency of echinocandins on the presence of nutrients is consistent with their killing activity being dependent on proliferating cells, whereas the weak metabolic dependency of amphotericin B, a pore former, is consistent with its ability to kill both proliferating and non-proliferating cells.

**Figure 6.**
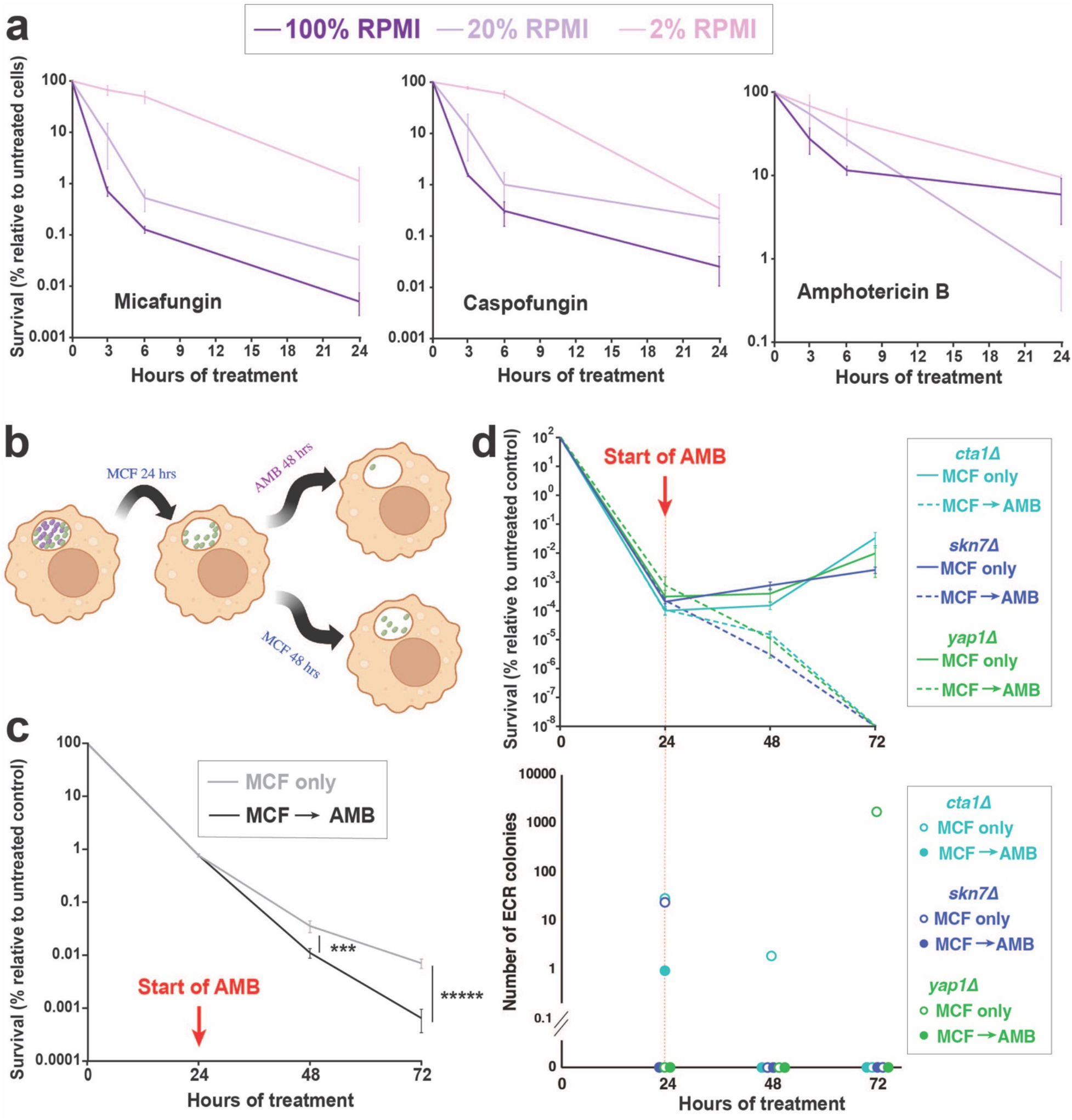
Amphotericin B efficiently kills echinocandin persisters. **a.** The metabolic dependencies of micafungin, caspofungin, and amphotericin B were assessed by incubating *C. glabrata* (CBS138) at various dilutions of RPMI (100%, 20%, and 2%). **a.** Micafungin and caspofungin killing efficiencies were much more affected by RPMI concentration than that of amphotericin B. **b.** ICG were treated with micafungin (0.125 μg/ml) for 24 hours, then treated with fresh RPMI containing either amphotericin B (2 μg/ml) or micafungin (0.125μg/ml), and survival rates were monitored for the next 48 hours. **c.** Following micafungin with amphotericin B significantly reduced the numbers of persister cells (* = p<0.03, ** = p<0.003, modified t-test). **d.** The same experiment was repeated with planktonically growing *cta1Δ, skn7Δ*, and *yap1Δ* mutants, and both survival and ECR colony frequency were assessed. Both survival and ECR frequencies were significantly reduced by following micafungin treatment with amphotericin B. ICG: intracellular *C. glabrata*, ECR: echinocandin-resistant, MCF: micafungin, AMB: amphotericin B.

Next, we asked whether amphotericin B could effectively kill ICG that had survived micafungin treatment. We treated ICG with micafungin (0.125 μg/ml) for 24 hours, followed by either replacing the medium with RPMI containing amphotericin B (2 μg/ml) or fresh RPMI containing micafungin (0.125 μg/ml), and survival was measured at 24- and 48-hour timepoints. Interestingly, our results showed that ICG treated with amphotericin B had a significantly lower survival rate compared to ICG treated with micafungin alone (Figure 6b).

Finally, we asked whether the amphotericin B-mediated increase in ICG killing was accompanied by reduced emergence of ECR colonies. We selected three ROS detoxification mutants showing a high ECR frequency following micafungin exposure (*cta1Δ, skn7Δ*, and *yap1Δ*), treated them with micafungin (0.125 μg/ml) for 24 hours to deplete drug-sensitive non-persisters, and then either replaced the medium with fresh RPMI containing amphotericin B (2 μg/ml) or fresh RPMI containing micafungin (0.125 μg/ml). ECR frequency was measured at 48- and 72-hour timepoints. Like the WT strain, the mutants had significantly higher survival when treated only with micafungin than when treated with micafungin followed by amphotericin B, and as expected, the higher survival was accompanied by higher ECR frequencies (Figure 6c), which could partly be explained by the fact that amphotericin B can indistinguishably kill both ECR and susceptible colonies. Collectively, these results indicate that alternating treatments with echinocandins and amphotericin B may be an effective strategy to eliminate intracellular *C. glabrata* persister reservoirs and decrease the frequency of emergence of ECR isolates.

## Discussion

*C. glabrata* is notable for its ability to proliferate inside macrophages (4, 55). Yet, the properties of ICG, particularly with respect to antifungal drug susceptibility, have not been examined. Our results identify macrophages as an important host reservoir of *C. glabrata* persistence and help explain how this fungal pathogen is able to rapidly generate antifungal drug-resistant mutations during treatment (9–14). We show that ICG cells exhibited hallmarks of bacterial persisters, such as increased survival in the presence of cidal drugs, a biphasic killing curve, lack of proliferation, lack of genetic mutations associated with mechanism specific resistance, reduced culturability, and the ability to reinfect macrophages. We also identify macrophage-generated ROS as an important inducer of *C. glabrata* drug tolerance and demonstrate that deleting of ROS detoxification genes greatly induces the emergence of ECR mutations. Finally, we show that intra-macrophage persister cells formed during echinocandin treatment can be eliminated by amphotericin B. Together, these results provide insights into how fungal pathogens persist in the host despite treatment with cidal antifungal drugs. These insights could be leveraged to develop more effective antifungal regimens, e.g., by alternating treatments with echinocandins with short treatments with amphotericin B, which would decrease the amphotericin B-associated toxicity while helping eliminate fungal persister cells and reducing the emergence of drug-resistant strains.

Our observation that ROS generated both *in-cellulo* and *in-vitro* increased the survival of *C. glabrata* upon exposure to a lethal concentration of micafungin is consistent with observations made using intracellular *Staphylococcus aureus* cells, where ROS likewise induced antibiotic tolerance (22, 23). How ROS induces drug tolerance and persistence is not well understood, but it has been shown that ROS can corrupt iron-sulfur containing enzymes involved in tricarboxylic acid (TCA) cycle (22, 23), which can result in lower respiration and ATP depletion (46, 60). Indeed, our observations indicate that ICG cells had significantly lower ATP levels than planktonic cells, which in turn may dampen the anabolic cellular processes requiring ATP, such as cell wall and membrane biosynthesis, and thus decrease the activities of antifungal drugs that target those cellular processes. Echinocandins’ killing activity may be more dependent on the metabolic/proliferative state of fungal cells because their target – the beta-glucan synthase enzyme – is specifically required by growing cells building and remodeling their cell walls. In contrast, amphotericin B can form pores in the membranes of both growing and non-growing cells, helping explain its weak dependence on the fungal metabolic status.

Interestingly, our previous RNAseq analysis of *C. glabrata* treated with echinocandins *in vitro* showed a strong increase in ROS levels but a concomitant downregulation of genes involved in ROS detoxification (28), suggesting that this ROS increase is a programmed change enacted by *C. glabrata* cells. Indeed, we previously found no evidence of ROS-based oxidative damage to DNA or lipids, suggesting that the ROS produced by *C. glabrata* during echinocandin treatment are not harmful to the cells. Consistent with this conclusion, here we show that deletion of genes involved in ROS detoxification did not reduce *C. glabrata* susceptibility to echinocandins *in vitro* or inside macrophages and that furthermore, deletion of these genes resulted in a significant increase in the number of ECR colonies. This result suggests that in echinocandin-treated cells ROS in addition to its many effector functions may act as a signal or a stimulant for generating mutations promoting drug resistance. The one intriguing exception to this pattern was the *icl1Δ* mutant that, despite containing increased ROS levels, did not produce any ECR colonies, which may indicate its involvement in multiple aspects of *fks* mutagenesis.

In bacterial literature, the term “tolerance” has emerged to indicate that the entire population is less affected by an antibiotic (e.g., by showing delayed killing dynamics), whereas “persistence” generally refers to a small subset of the population that can survive long-term antibiotic exposure (17). Both persistence and tolerance lack drug target alterations, and both reproducibly give rise to progenies that exhibit survival/growth phenotypes similar to the parental strains. In general, in medical mycology the terms “tolerance” and “persistence” have been used somewhat interchangeably (28, 61, 62), and this has been further complicated by the usage of the term “tolerance” for both cidal and static drugs. For the former, tolerance refers to survival in the presence of supra-MIC drug concentrations, whereas for the latter, tolerance refers to incomplete growth inhibition. In this study we focused on cidal drugs and characterized a sub-population of cells able to withstand their killing activity, which we term “persisters” according to bacterial nomenclature. Given that both tolerance and persistence are potentially of clinical importance and that mechanisms underpinning them could be different, appreciation of their differences can inspire detailed molecular studies to better comprehend the biology of drug-refractory fungal infections.

Altogether our results indicated that macrophages constitute a permissive reservoir for development of drug persister *C. glabrata* cells, which facilitate the emergence of stable drug resistance. Finally, amphotericin B was featured as a promising therapeutic potential to significantly reduce the repertoire of micafungin persister and resistant *C. glabrata* cells.

## Methods

### *C. glabrata* strains and growth conditions

We used 8 *C. glabrata* isolates, including 7 clinical isolates and a type-strain CBS138 (ATCC2001), which all had the same MIC values, but belonged to different sequence types and various geographical locations (Table S1). The deletant mutants were all derived from the reference strain CBS138. *C. glabrata* cells were grown on yeast-peptone-dextrose (YPD) agar plates overnight at 37°C and the initial inoculum used for the macrophage infection or evaluation of in-vitro stresses were incubated in YPD broth overnight at 37°C.

### THP1 macrophages and growth conditions

Human acute monocyte leukemia cell line-derived macrophages (THP1; ATCC; Manassas, VA) was used to assess the phagocytosis survival of our *C. glabrata* isolates. RPMI 1640 (Gibco, Fisher Scientific, USA) supplemented with 1% penicillin-streptomycin (Gibco, Fisher Scientific, USA) and 10% heat-inactivated HFBS (Gibco, Fisher Scientific, USA) was used to grow the THP1 cells. One million THP1 cells treated with 100 nM phorbol 12-myrisate 13-acetate (PMA, Sigma) were seeded into 24-well plates and incubated at 37°C in 5% CO_2_ for 48 hrs to induce attachment and differentiation into active macrophages. At the day of infection macrophages were washed with PBS, fresh RPMI was added, and *C. glabrata* cells were exposed to macrophages. Subsequently, the plates were centrifuged (200g, 1 minute), and incubated at 37°C in 5% CO2 for 3 hrs. In the next step, the supernatant RMPI was removed, and the pellets were five times washed with phosphate buffered saline (PBS) solution. Macrophage lysis was performed by adding 1 ml cold water (kept at 4°C) and 100 μl of the lysed macrophages were serially diluted, plated on YPD agar, and incubated at 37°C for up to 72 hrs.

### Sequencing of HS1 and HS2 of *FKS1* and *FKS2*

HS regions of ECR colonies emerging on YPD plates containing micafungin were amplified and sequenced using primers and conditions described elsewhere. The WT sequences of *FKS1* and *FKS2* from ATCC2001 were used as our control (50).

### Measuring killing dynamic and recovery of ECR colonies following exposure to micafungin

PMA treated THP1 macrophages were infected with *C. glabrata* isolates belonging to different STs with the MOI of 10/1 (10 *C. glabrata* cells/1 macrophage), which served as our treated groups. Since *C. glabrata* can dramatically replicate inside the macrophages if not treated with micafungin and if incubated for a long time, the THP1 macrophages of the untreated groups were infected with the MOI of 1/10. Three hrs postexposure, macrophages were extensively washed with PBS and the RPMI containing 0.125μg/ml of micafungin was used for the treated group, while only fresh RPMI was added to the control group. Simultaneously, respective *C. glabrata* isolates were incubated in RPMI containing micafungin and untreated groups were grown in drug-free RPMI. At each time-point 100 μl of the lysate was plated on YPD agar and the rest were centrifuged, the supernatant was decanted, 200μl of PBS was added, and resuspended cells were transferred to YPD plates containing 0.125μg/ml of micafungin (from 24-hrs onward). We chose 0.125μg/ml of micafungin, since it could detect all the ECR *C. glabrata* isolates harboring various clinically relevant mutations and since the incubation times for the detection of ECR colonies varied depending on mutation type and ECR cell number (we used serial dilution), we extended the incubation time (37°C) to 7 days to ensure that we could capture all ECR colonies regardless of mutation type and cell number. All treated and untreated groups from both macrophages and RPMI arms were plated on YPD agar containing micafungin to monitor if any ECR colonies could emerge from the untreated groups. The dynamic of killing was measured up to 120 hrs and survival rate of the treated groups were normalized against untreated group of respective time-points. Of note, since the number of *C. glabrata* cells of treated groups were 100 times higher than the untreated counterparts, this dilution factor was considered in our normalization.

### Measuring intracellular concentration of micafungin

THP-1 monocytes grown in supplemented RPMI 1640, were seeded into 96-well tissue culture-treated plates at 5x10^4^ cells / well. THP-1 monocytes were differentiated overnight to macrophages with 100 nM phorbol 12-myristate 13-acetate. Culture medium was replaced with fresh medium containing micafungin, rifampicin, moxifloxacin or clofazimine. Micafungin was assayed at 3.14 μM and drug controls were assayed at 5 μM. Each drug was tested in triplicate wells. After 0.5, 3 and 24 h incubation at 37°C, the cells were washed twice with cold PBS to remove extracellular drug. Cells were extracted with 60% DMSO for 1 h at 37°C. Micafungin concentration in each sample was analyzed by LC-MS/MS, and normalized by the number of cells per well and the average THP-1 cellular volume to calculate intracellular concentrations (63, 64). Intracellular drug accumulation is expressed as the ratio between the intracellular concentration and extracellular concentration (IC/EC).

### LC-MS/MS

Neat 500μM micafungin solution was serially diluted in 50/50 acetonitrile (ACN)/water to create neat standard curves and quality control spiking solutions. 10 μL of neat spiking solutions were added to 90 μL of drug-free macrophage lysate to create standard and QC samples. 10 μL of control, standard, or study sample lysate were added to 100 μL of a 50:50 acetonitrile: methanol protein precipitation solvent mix containing 10 ng/mL of the internal standard verapamil to extract micafungin. Extracts were vortexed for 5 minutes and centrifuged at 4000 RPM for 5 minutes. 75 μL of supernatant was transferred for LC-MS/MS analysis and diluted with 75 μL of Milli-Q deionized water. LC-MS/MS analysis was performed on a Sciex Applied Biosystems Qtrap 6500+ triple-quadrupole mass spectrometer coupled to a Shimadzu Nexera X2 UHPLC system to quantify each drug concentrations in each sample. Chromatography was performed on an Agilent SB-C8 (2.1x30 mm; particle size, 3.5 μm) using a reverse phase gradient. Milli-Q deionized water with 0.1% formic acid was used for the aqueous mobile phase and 0.1% formic acid in acetonitrile for the organic mobile phase. Multiple-reaction monitoring of parent/daughter transitions in electrospray positive-ionization mode was used to quantify all analytes. The MRM transitions of 455.40/165.20, 1270.40/1190.40, 823.5/791.6, 402.2/358.0, 473.2/431.2 were used for verapamil, micafungin, rifampicin, moxifloxacin and clofazimine, respectively. Data processing was performed using Analyst software (version 1.6.3; Applied Biosystems Sciex).

### Measuring the survival of planktonic and ICG cells after exposure to cidal antifungal drugs

THP1 macrophages were infected with MOI of 10/1 and after 3 hrs they were washed extensively to remove non-adherent cells. Macrophages were lysed using ice-cold water followed by vigorous pipetting and *C. glabrata* cells were collected using centrifugation. One ml of RPMI 1640 containing micafungin (0.06μg/ml), caspofungin (0.125 μg/ml), and amphotericin B (2μg/ml) were added to collected cells and cell suspensions were incubated at 37°C for 1 hour. Prior and 1 hour after incubation suspensions were plated on YPD plates, which were incubated at 37°C for 1 day. The CFU of treated cells were normalized against those prior to exposure and the values were presented as percentage.

### FITC and AF-647 ConA staining

PBS washed initial inoculum of *C. glabrata* cells were stained with 200μg/ml of FITC (Millipore Sigma) in carbonate buffer (0.1M Na_2_CO_3_, 0.15M NaCl, pH= 9.3) and incubated at 37°C, followed by 3 times PBS washing. The THP1 macrophages were infected with MOI of 10/1 and after 3-, 6-, 24-, and 48-hrs pst with micafungin (0.125μg/ml), the *C. glabrata* cells were released from macrophages and counterstained with 50μg/ml AF-647-ConA (Millipore Sigma) in PBS buffer+2% BSA. The pellets were washed 3 times with PBS and subjected to flow cytometry (BD Biosciences). Double-stained yeasts were mother cells, while yeast cells only stained with AF-647 ConA represented replicated cells.

### PI staining

*C. glabrata* cells released from macrophages were treated PBS containing 10μg/ml of propidium iodide (PI, Millipore Sigma) and analyzed by FACS (Melody instrument, BD Biosciences). PI^-^ cells were collected and used for ATP determination.

### ATP measurements

The yeast cells were released from macrophages by adding 0.2% Tritron 100X and they were collected by centrifugation. The pellets were rewashed with ice-cold water, followed by vigorous vortexing and recollecting by centrifugation. *C. glabrata* cells were treated with 500μl of Y1 buffer (1M of sorbitol and 0.1M EDTA) containing 100 units of lyticase and the pellets were incubated at 37°C for 30 minutes. Subsequently, the spheroplasts were collected by centrifugation, the supernatant was decanted, and 200μl of lysis buffer (10mM Tris, 1mM EDTA, 0.01% triton, and 100mM NaCl) was added to each tube, followed by incubation at 80°C for 10 minutes and then subjected to bead beating for 2 minutes at the highest speed. 20μl of this lysate was subjected to 180μl of luciferase reaction (ThermoFisher Scientific) and luminescence rate was measured using a plate reader.

### Mouse systemic infections

Six-week-old CD-1 female mice were used for in-vivo systemic infections. Four days prior to infection, mice were immunosuppressed using 150mg/kg of cyclophosphamide and immunosuppression with 100mg/kg of cyclophosphamide was continued every 3 days afterward. At the day of infection (day 0), 50μl of cell suspension containing 5x10^7^ cell was administered intravenously. Mice were grouped into two arms (*n*=24), treated with 5mg/kg of caspofungin starting 4 hrs after infection and continued until the end of experiment (12 mice), and the second arm served as control, which included mice treated with PBS only (12 mice). Four mice per each group were euthanized and sacrificed at day 1, 4, and 7, spleen and kidneys were harvested and homogenized, and 100μl of each homogenate was plated on YPD plate and incubated for 24-48 hrs at 37°C. The number of colonies from each organ of treated mice were normalized against the respective organ of untreated group and data presented as percentage.

### Generation of *C. glabrata* knock-out mutants

The following genes were subjected to deletion, catalase 1 (*CAT1*, CAGL0K10868g), glutathione oxidoreductase (*GRX2*, CAGL0K05813g), manganese superoxide dismutase (*SOD2*, CAGL0E04356g), and three transcription factors playing role in oxidative stress responses, namely *SKN7* (CAGL0F09097g), *MSN4* (CAGL0M13189g), and *YAP1* (CAGL0H04631g), and *ICL1* (CAGL0L09273g). Knock-out mutants were created in house using a previously described protocol (65), in which the open reading frame of the gene of interest was replaced by nourseothricin (NAT) resistance cassette. The knock-out construct was generated by using Ultramer primers (~80-100 bps) containing homology regions with NAT and with regions flanking GOIs. Competent cells were created by log-phase grown *C. glabrata* cells using Frozen-EZ Yeast Transformation Kit (Zymo Research) and transformation followed an electroporation-based protocol described previously (65). The colonies growing on YPD plates containing NAT were subjected to PCR and sequencing using diagnostic primers listed in Supplementary Table 3 (the primers used in the current study were manufactured by Integrated DNA Technologies and Sanger Sequencing carried out by Genewiz). Two independent knock-out mutants were used for each experiment.

### Measuring metabolic dependency of antifungal drugs

Overnight grown *C. glabrata* cells (CBS138) were PBS washed thrice, enumerated, and 100μl of 10^8^ cells were added to 1 ml of various RPMI concentrations containing micafungin or caspofungin or amphotericin B. Desired concentrations of RPMI were simply made by addition of sterilized demi water to 100% RPMI. The survival of *C. glabrata* cells incubated in various 100%-, 20%-, and 2%-RPMI containing micafungin (0.125μg/ml) or caspofungin (0.25μg/ml) or amphotericin B (2μg/ml) was assessed at 3-, 6-, and 24-hrs pst (totaling 9 conditions for each time-point). *C. glabrata* cells incubated in respective RPMI concentrations lacking any antifungal drugs were considered as control. The survival rate of drug exposed *C. glabrata* cells were normalized against drug-free controls, which was presented by percentage. A higher metabolic dependency was defined when the lethality of a given antifungal drug significantly decreased at lower concentrations of RPMI.

### Measuring the impact of amphotericin B or micafungin treatment on ICG survival rate

ICG cells (CBS138) treated with micafungin (0.125μg/ml) for 24 hrs were exposed to 1ml of fresh RPMI containing either amphotericin B (2μg/ml) or micafungin (0.125μg/ml) and the survival rate was assessed 24 and 48 hrs later. The survival rates of treated ICG were normalized against the counterparts treated with drug-free RPMI and the data were presented by percentage.

### Measuring the impact of amphotericin B or micafungin treatment on ECR rate

Overnight grown knockout mutants, *cta1Δ, yap1Δ*, and *skn7Δ*, were PBS washed twice, enumerated, and 100μl of 10^8^ cells were added to 1 ml of 100% RPMI containing micafungin (0.125μg/ml) and incubated for 24 hrs. After 24 hrs the *C. glabrata* cells were collected, washed with PBS, and either treated with fresh RPMI containing micafungin (0.125μg/ml) or amphotericin B (2μg/ml) and survival rate and mutation frequency were assessed 24 and 48 hrs later. The respective *C. glabrata* mutants grown in drug-free RPMI were used as control.

### ROS measurement

ROS was measured using dihydroethidium (DHE) (6.5 μg/ml; ThermoFisher), which detects superoxide levels (28, 66, 67). *C. glabrata* isolates lacking *CTA1, GRX2, ICL1, MSN4, SKN7, SOD2*, and *YAP1* and the parental WT strain of CBS138 were incubated in RPMI (Gibco™ RPMI 1640 Medium) containing 0.125μg/ml of micafungin for 2 hrs. CBS138 incubated in drug-free served as our negative control. Since our previous study using the same dyes showed the lack of ROS detection from ECR isolates, we did not include ECR isolates in this experiment (28). After each time-point, *C. glabrata* cells, at least in three biological replicates, were collected, washed with prewarmed PBS, and stained with DHE in PBS for 30-40 minutes at 37°C. Subsequently, the stained cells were collected, 1 ml of prewarmed PBS was added and subjected to flow cytometry and the data obtained were analyzed by FlowJo software v10.6.1 (BD Biosciences).

### Antifungal susceptibility testing (AFST)

The broth microdilution protocol of the CLSI M27-A3 was followed. AFST included the following antifungal drugs, fluconazole (Pfizer), amphotericin B (Sigma-Aldrich), micafungin (Astellas Pharma), and anidulafungin (Pfizer). Plates were incubated at 37°C for 24 hrs, and the MIC50 data (50% growth reduction compared to controls without drug) were determined visually. Each experiment included at least three biological replicates.

