## Supplementary figures and images for "Macrophage internalization creates a multidrug-tolerant fungal persister population, providing a permissive reservoir for the emergence of drug resistance"

### Supplementary Figure 1

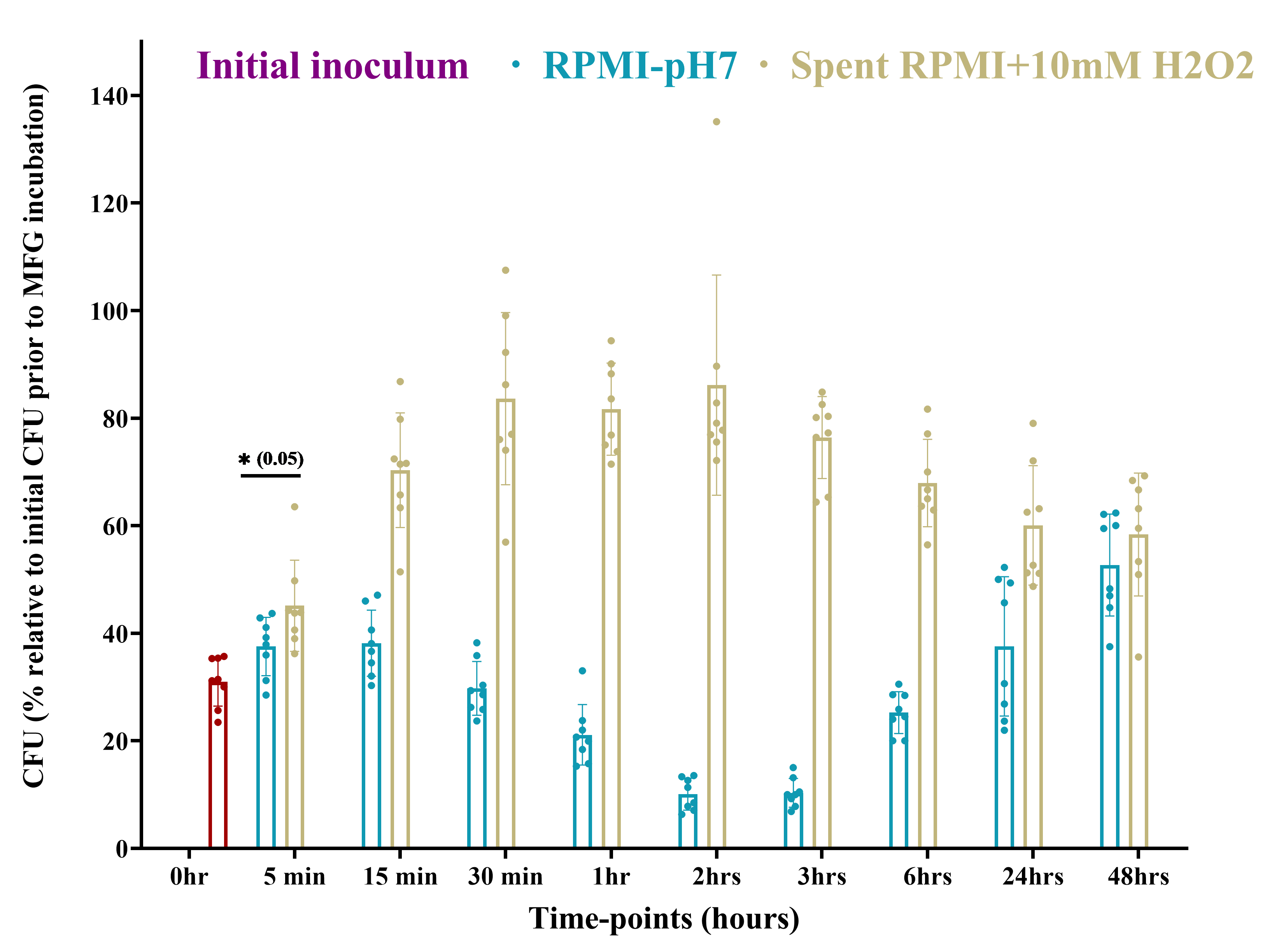
